# Mitigation of the age-dependent decline in mitochondrial genome integrity

**DOI:** 10.1101/2022.01.18.476860

**Authors:** Pei-I Tsai, Ekaterina Korotkevich, Patrick H. O’Farrell

## Abstract

Unknown processes promote accumulation of mitochondrial DNA mutations during aging. Accumulation of defective mitochondrial genomes is thought to promote progression of heteroplasmic mitochondrial diseases and degenerative changes with natural aging. We used a heteroplasmic *Drosophila* model to test 1) whether purifying selection acts to limit the abundance of deleterious mutations during development and aging, 2) whether quality control pathways contribute to purifying selection, 3) whether activation of quality control can mitigate accumulation of deleterious mutations, and 4) whether improved quality control improves healthspan. We show that purifying selection operates during development and growth, but is ineffective during aging. Genetic manipulations suggest that a quality control process known to enforce purifying selection during oogenesis also suppresses accumulation of a deleterious mutation during growth and development. Flies with nuclear genotypes that enhanced purifying selection sustained higher genome quality, retained more vigorous climbing activity and lost fewer dopaminergic neurons. Pharmacological enhancement of quality control produced similar benefits. Importantly, similar pharmacological treatment of aged mice reversed age-associated accumulation of a deleterious mtDNA mutation. Our findings reveal dynamic maintenance of mitochondrial genome fitness, and reduction in the effectiveness of purifying selection during life. Importantly, we describe interventions that mitigate and even reverse age-associated genome degeneration in flies and in mice. Furthermore, mitigation of genome degeneration improved wellbeing in a *Drosophila* model of heteroplasmic mitochondrial disease.

**Significance Statement:** In contrast to the orderly segregation of nuclear DNA, mitochondrial genomes compete for replication and segregation. The abundance of mutants that emerge in a cell depends on their success in this competition. We show that quality control mechanisms put deleterious mutations at a disadvantage, but these mechanisms become ineffective during aging. The resulting rise in mutant genomes, which compromises vigor, can be suppressed using genetic backgrounds that enhance quality control. Feeding kinetin to adult *Drosophila* or mice also reduced the load of mutant mitochondrial genomes. This pharmacological reversal of the age associated deterioration of mtDNA quality suggests possible therapies to alleviate mitochondrial diseases and normal aging.

## Introduction

Unlike nuclear genotype, which is largely stable during one’s lifetime, when genetically distinct mitochondrial genomes co-reside (heteroplasmy), their relative proportions shift during growth, development, and aging. This shift is not random. Mutant mtDNA variants accumulate during aging and in the progression of some mitochondrial diseases (1–5). Stereotyped changes in abundance of particular alleles in different tissues in human and mouse indicate that selective forces favor different mitochondrial genomes (3, 6, 7). However, despite the importance of mitochondrial function to health and wellbeing, we have limited understanding of the processes underlying accumulation of mitochondrial mutations with age.

The nuclear genome encodes mechanisms of quality control that survey the function of mitochondria and eliminate or compromise the proliferation of defective mitochondria (8, 9). Two described mechanisms use PINK1, the product of a gene discovered as one of the causes of early onset familial Parkinson’s disease, as a sensor of mitochondrial function. PINK1 accumulates on the surface of mitochondria having a reduced membrane potential and its kinase activity in this location signals several downstream events (10–15). In one pathway defined largely in a cell culture model, PINK1 activates PARKIN, the product of another Parkinson’s disease gene, which then triggers elimination of compromised mitochondria by mitophagy (16). In a second pathway, acting in the *Drosophila* female germline (17, 18), PINK1 acts in a PARKIN independent pathway that targets a protein called Larp to inhibit its role in promoting biogenesis of the mitochondria (18) (SI Appendix, Fig. S2).

Since deleterious mtDNA mutations compromise electron transport of mitochondria, it seems that quality control would put genomes carrying such mutations at a disadvantage creating a purifying selection that leads to their elimination. However, this outcome is far from certain. Various factors such as the sharing of gene products among mitochondria as a result of dynamic fission and fusion could mask the consequences of heteroplasmic mutations shielding them from quality control. Indeed, a number of studies suggest a contrast between the germline, which exhibits purifying selection, and adult somatic tissues, which often show accumulation of mutations. Studies in *Drosophila* show that purifying selection acting in the female germline eliminates deleterious mutations in a few generations (18–21). Genetic dissection revealed that this purifying selection depends on the PINK1/LARP pathway of quality control (18). On the other hand, an elegant study that induced heteroplasmic deletions in the flight muscle of the adult fly detected no substantial indications of purifying selection unless additional stressors were introduced (22). Furthermore, a detailed study in *Drosophila* carrying a proofreading defective mitochondrial DNA polymerase (mutator line) showed that adult flies accumulate a spectrum of mutations biased toward deleterious mutations (23). Since deleterious mutations would potentially be removed by purifying selection, this finding argued either that it was not operating or was opposed by a stronger selection favoring deleterious mutations (23). Similarly, studies in mouse suggest a discordance between germline and soma. When mutations in the mitochondrial genome that were introduced in a mutator line were passed through subsequent generations in a wildtype background, there was selective elimination of deleterious mutations, a strong signal of purifying selection (7). In contrast, zygotically accumulated mutations in the mutator mouse exhibited high levels of deleterious mutations, suggesting a lack of purifying selection in the soma. Furthermore, Parkin mutant mice did not show a significant increase in mutations in adult wild-type or mutator mice, suggesting that Parkin dependent quality control does not contribute to purifying selection (16). While these studies suggest major changes in the efficiency of purifying selection, other studies have suggested continued purifying selection in some circumstances. For example, adult human T-cells of mitochondrial disease patients show exceptionally low heteroplasmy levels suggesting cell-type specific action of purifying selection (24). We have sought to measure purifying selection and to understand the nature of quality control in the soma during growth, development, and aging using an experimental model developed in *Drosophila*.

A previously described heteroplasmic line of *D. melanogaster* carries a wild-type mitochondrial genome (*Yak*-mt) from another species, *D. yakuba*, and a *D. melanogaster* genome (*Mel*-mt^ts^) crippled by a temperature sensitive mutation in cytochrome oxidase subunit 1 (*mt:CoI*^*T300I*^)(Fig. 1*A* and (25)). The *Mel*-mt^ts^ genome has an intrinsic advantage in replication. At permissive temperature it gradually displaces the *Yak*-mt genome resulting in the loss of the *D. yakuba* genome in a few generations (25). At 29°C, a temperature at which the *CoI*^*T3000I*^ mutant cannot support viability (26), purifying selection counters the replicative advantage of the mutant *Mel*-mt^ts^ genome preventing it from taking over (25). qPCR gives a measure of the ratio *Yak*-mt/total-mt. The difference in this ratio between permissive and restrictive temperatures allows us to assess the impact of a functional disparity on competition between the two genomes and provides a measure of purifying selection.

**Figure 1.**
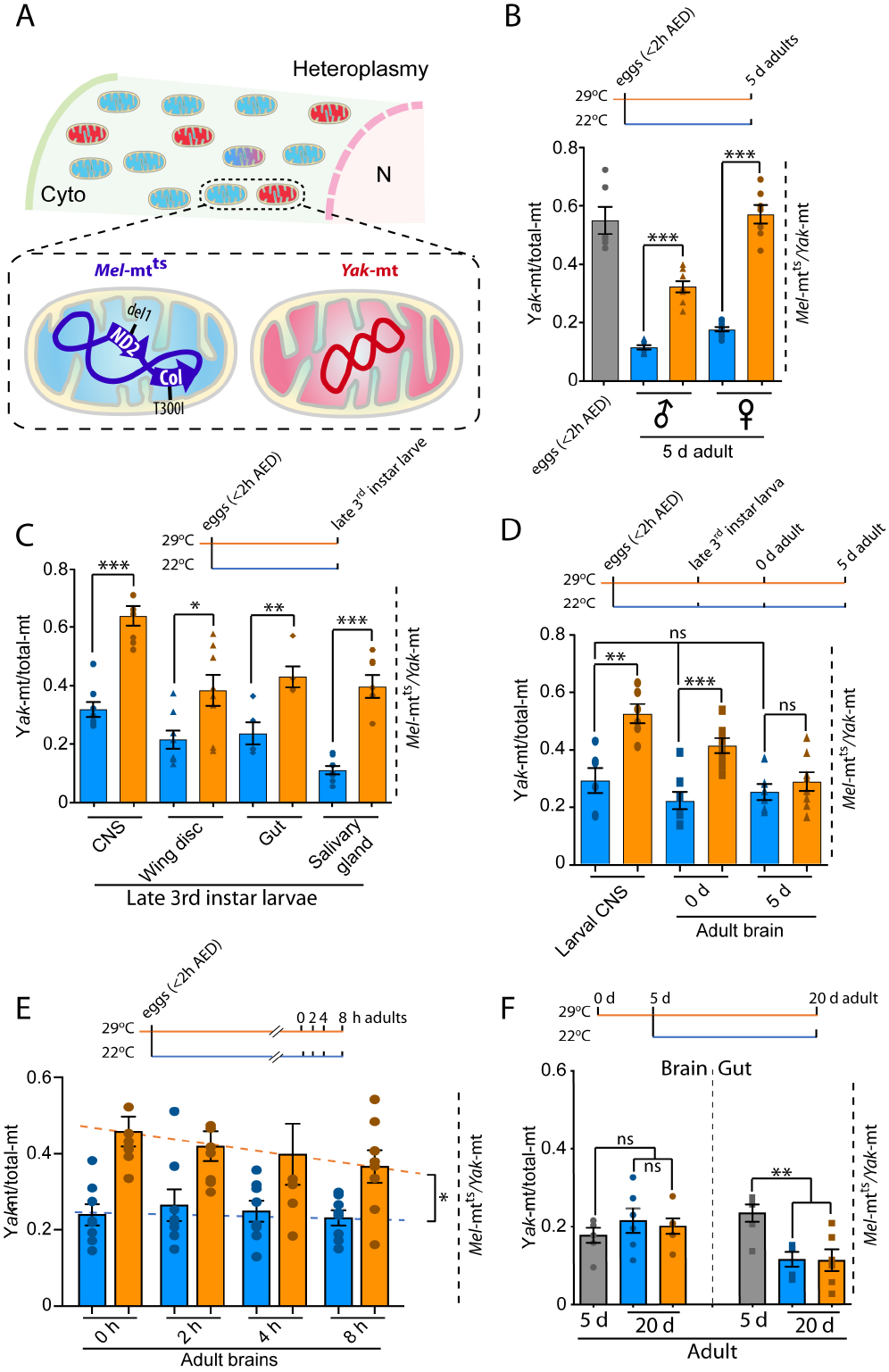
Quality control modulates the ratio of heteroplasmic mitochondrial genomes during development. **(*A*)** Heteroplasmy for schematized *Yak-*mt and *Mel-*mt^ts^ genomes was established by transferring cytoplasm of *D. yakuba* embryos into *D. melanogaster* embryos carrying the doubly mutant genome *mt:ND2*^*del*^ *+ mt:Col*^*T300I*^ (*Mel-* mt^t*s*^). **(*B*)** The proportion of *Yak-*mt (*Yak-*mt/total-mt) following development from egg to adult shows action of quality control. Eggs (2 h collection at 29°C) were assayed (grey bar) or allowed to develop to 5d post eclosion at 22°C (blue) or 29°C (amber). Adult females have a large contribution from oocyte mtDNA. **(*C*)** Quality control operates in multiple tissues with different effectiveness. The blue and amber bars (22°C or 29°C, respectively) show the proportion of *Yak-mt* in different tissues of late 3^rd^ instar larvae. **(*D*)** The impact of purifying selection declined with age in the CNS. **(*E*)** The decline in relative abundance of wild-type *Yak*-mt following eclosion was fast. **(*F*)** The signature of quality control is absent during maturation of adults. *Yak*-mt/total-mt ratios from gut and brain taken from 5-day (grey bars) or 20-day adults aged at 22°C or 29°C (blue and amber bars, respectively). Here and below, * = *p* < 0.05, ** = *p* < 0.01, and * = *p* < 0.001 by one-way ANOVA/Tukey’s multiple comparison test. Data represent 8 independent biological repeats, with each repeat being an average of ratios assessed in three samples of eggs or adults. For tissues, data represents tissues dissected from 8 individuals. AED, after egg deposition. In **(*E*)**, slopes differ * = *p* < 0.05 by liner regression.

Past work using this heteroplasmic line focused on changes in the relative abundance of the two genomes from one generation to the next, and uncovered the action of purifying selection during oogenesis (18–21, 25, 27). It is notable that this selection, depends on quality control, occurs by competition between mitochondria within the oocyte, and does not involve selection for organismal fitness (18, 21). Changes in the ratio of *Yak*-mt/total-mt during the lifetime of the fly suggested that maintenance of this ratio is dynamic (25). Here, we have examined changes in the relative abundance of the two genomes in the soma. Shifts in the ratio of *Yak*-mt/total-mt provide a measure of the effectiveness of purifying selection during the life of the fly and a means of assessing whether purifying selection can be genetically or pharmacologically modified. We find evidence that purifying selection is active in the soma during growth and development. The influence of mutations in quality control genes suggests that somatic mechanisms of quality control overlap those operating in the germline. However, the effectiveness of purifying selection declines with age and has no obvious impact in tested tissues beyond 5 days after eclosion of adult flies. Importantly, we find that quality control can be stimulated in the adult by genetic alterations or by feeding kinetin and that such measures forestall mutational accumulation and aging phenotypes in *Drosophila* and can reverse mutational accumulation in aged mice.

## Results

### Somatic selection of mitochondrial genomes during development and aging

To assess the action of selection in somatic tissues during development, we collected eggs from heteroplasmic flies at 29°C, allowed development to young adulthood at either 22°C or 29°C, and measured the ratios of mitochondrial genomes in whole flies (Fig. 1B). For analysis of events in the soma, we focus on males, because high ovarian mtDNA levels result in a large germline influence in females (25). At 22°C, the ratio of *Yak*-mtDNA to total-mtDNA declined dramatically from egg to adult, representing the outcome of competition when both genomes were functional. At 29°C, *Yak*-mt/total-mt was maintained at a higher level than in flies raised at 22°C. This difference reflects a response to the functional disparity between the genomes, and shows selection against the detrimental mutation. However, the *Yak*-mt/total-mt ratio in the 5-day adult males was lower than in freshly laid eggs, suggesting that this selection is weaker during growth to adulthood than it was in the female germline (Fig. 1B).

To assess the influence of a functional disparity during earlier development, we examined larval tissues after development at 22 or 29°C. Again, the *Yak*-mt/total ratio declined at 22°C. The extent of the decline was different in the different tissues. Although, the basis for this difference is not known, it might be attributed to tissue-specific influences on the replicative abilities of the two different genomes (3). However, regardless of the cause, the *Yak*-mt/total ratio at 22°C serves as a control for change induced when *mt:CoI* is inactivated by increasing the temperature to 29°C. Total mtDNA levels were similar at both temperatures (*SI Appendix*, Fig. S1), suggesting that copy-number is not responsive to the difference in *mt:CoI* function. In contrast, the ratio of *Yak*-mt to total-mt was higher at 29°C in all tissues examined (Fig. 1*C*). We conclude that some form of purifying selection contributes to mitochondrial genome quality in diverse larval tissues during development to the late larval stage.

Despite early purifying selection in larval stages and maintenance at 29°C, the *Yak*-mt/total-mt ratio in the CNS/brain declined during continued development (Fig. 1D). Indeed, 5 days after eclosion from the pupal case (5-day adult) the ratio was nearly at the level seen in the brains of flies that were raised the entire time at 22°C (Fig. 1D). Measuring *Yak*-mt/total-mt every 2 hours after eclosion revealed a rapid decline of the ratio (Fig. 1E). Thus, the transition to adulthood is accompanied by a rapid shift in the ratio of mitochondrial genomes that erases the benefits of earlier purifying selection in the brain. Purifying selection must decline before the decline has its consequence on the *Yak*-mt/total-mt ratio. Thus, the rapidity of the post-eclosion readjustment of *Yak*-mt/total-mt reveals a dynamic requirement for purifying selection.

Aging of adult flies from 5 to 20 days did not result in further decline of *Yak*-mt/total-mt in brain at either temperature. In contrast, the *Yak*-mt/total-mt fell further in the highly proliferative gut. However, it is notable that this decrease in the gut is the same at both temperatures. The temperature independence of *Yak*-mt/total-mt during aging indicates a lack of effective purifying selection against the detrimental mutation after 5 days of adulthood.

### Nuclear genes modulate purifying selection of mitochondrial genomes in the soma

Since PINK1 acts as a sensor in quality control pathways, we tested its influence on purifying selection in the soma. A loss-of-function mutation in the X chromosomal *Pink1* locus dramatically decreased *Yak*-mt/total-mt in 5-day adult male flies raised at 29°C (Fig. 2A). This decline was significantly rescued by an autosomal copy of wild-type PINK1. We conclude that *Pink1* contributes to somatic purifying selection in flies.

**Figure 2.**
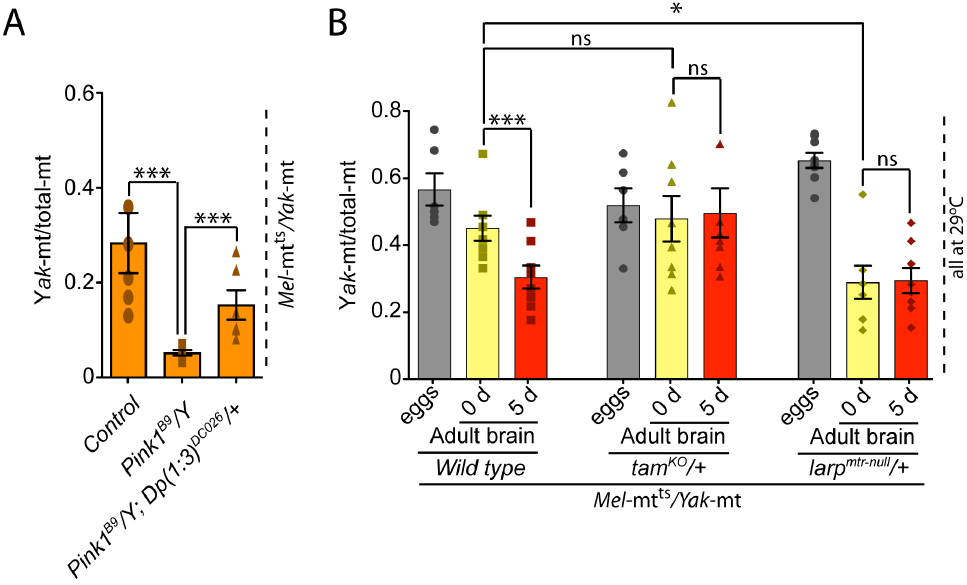
The nuclear genotype impacts the maintenance of mitochondrial genome quality in adult brains. **(*A*)** Loss-of-function of *Pink1* leads to decline in *Yak-*mt/total-mt indicating that PINK1 helps sustain high levels of the functional genome. Data show genome ratios for 5-day old flies of the indicated genotype raised at 29°C. The *Pink1*^*B9*^/*Y* flies lack Pink1 function, while the *Pink1*^*B9*^/*Y*; *Dp(1:3)*^*DC026*^/*+* flies carry a rescuing autosomal duplication of the normally X chromosomal *PINK1* gene. The PINK1 deficient flies and rescued flies are sibs from the same cross and therefore have the same pool of maternally contributed genomes. Note also, that based on the *Drosophila* version of dosage compensation, the single X chromosomal copy of PINK1 in the control is expected to have twice the activity of the rescuing autosomal copy. **(*B*)** The influence of nuclear genotype on mitochondrial genome quality of adult brains. Bars show the ratio of *Yak-*mt/total-mt in eggs (grey), and in brains of newly eclosed (0 d: yellow) or 5-d old (red) adults when the indicated genotypes are raised at 29°C. Data reflect the consequences of zygotic dose-reduction of the tested gene.

To assess quality control events downstream of *Pink1*, we reduced the gene dose of *larp*, a mediator of *Pink1* quality control in the germline (*SI Appendix*, Fig. S2) (18). Although changes in gene dose usually produce subtle or no phenotype, we reasoned that the decline in the *Yak*-mt/total-mt ratio in the brains during the transition from pupa to adulthood might provide an especially sensitive point to monitor the impact of quality control in somatic tissues. Heterozygosity for *larp* compromised purifying selection so that the brains of newly eclosed flies had much reduced *Yak*-mt/total-mt (Fig. 2B). Additionally, a genome-wide screen showed that reduction in the dose of the gene *tamas* (*tam*), which encodes the mitochondrial DNA polymerase catalytic subunit (POLγA), enhanced elimination of mitochondrial genomes with deleterious mutations during oogenesis (27). When we reduced the dose of *tam* during growth and development, a higher *Yak*-mt/total-mt persisted in brains indicating stronger purifying selection (Fig. 2B, *SI Appendix*, Fig. S3, S4, S5). These findings show parallels between germline and somatic purifying selection and implicate a common quality control pathway in the processes.

Adult *Yak*-mt/total-mt heteroplasmic flies are lethargic and short lived (about 20 days) at 29°C. Since heteroplasmic Pink1 mutants had a reduced mitochondrial genome quality, we examined the influence of *Pink1* on the health of heteroplasmic flies. *Pink1* mutant flies that are otherwise wild type are viable at 22 and 29°C. In contrast, while *Pink1* flies heteroplasmic for *Yak*-mt and *Mel*-mt^ts^ survive at 22°C, they die during the first few days of adulthood at 29°C. This finding shows that PINK1 contributes to survival of heteroplasmic flies when function of *Mel*-mt^ts^ is compromised. The death is likely the consequence of the much-reduced abundance of the functional mitochondrial genome (Fig. 2A).

To assess possible benefits of increased *Yak-*mt/total-mt, we examined the consequence of reducing the dose of *tam* (*tam* heterozygosity). We measured the *Yak*-mt/total-mt ratio in control and *tam* heterozygous flies as they aged, and assessed their vigor using a climbing assay. Compared to controls, heterozygous (*tam*^*KO*^/+) flies sustained a higher level of *Yak*-mt/total-mt for 14 days into adulthood (Fig 3A). They also showed more vigorous climbing at least until 11 days after eclosion (Fig. 3B). Neuronal cell loss has been associated with aging-associated degenerative conditions. To test for such an association, we tested the PPL1 cluster of dopaminergic (DA) neurons for cell loss in control and in *tam*^*KO*^ heterozygous flies. Normal flies exhibit an invariant number of PPL1 DA neurons throughout adult life (28). In contrast, DA neuron number dropped between 5 and 20 days of adulthood in the heteroplasmic flies. This decline was not detected in the *tam*^*KO*^/+ heteroplasmic line (Fig 3D). The phenotypic benefits associated with maintenance of a higher proportion of functional genomes suggests that mitochondrial genome quality impacts age-associated degeneration.

**Figure 3.**
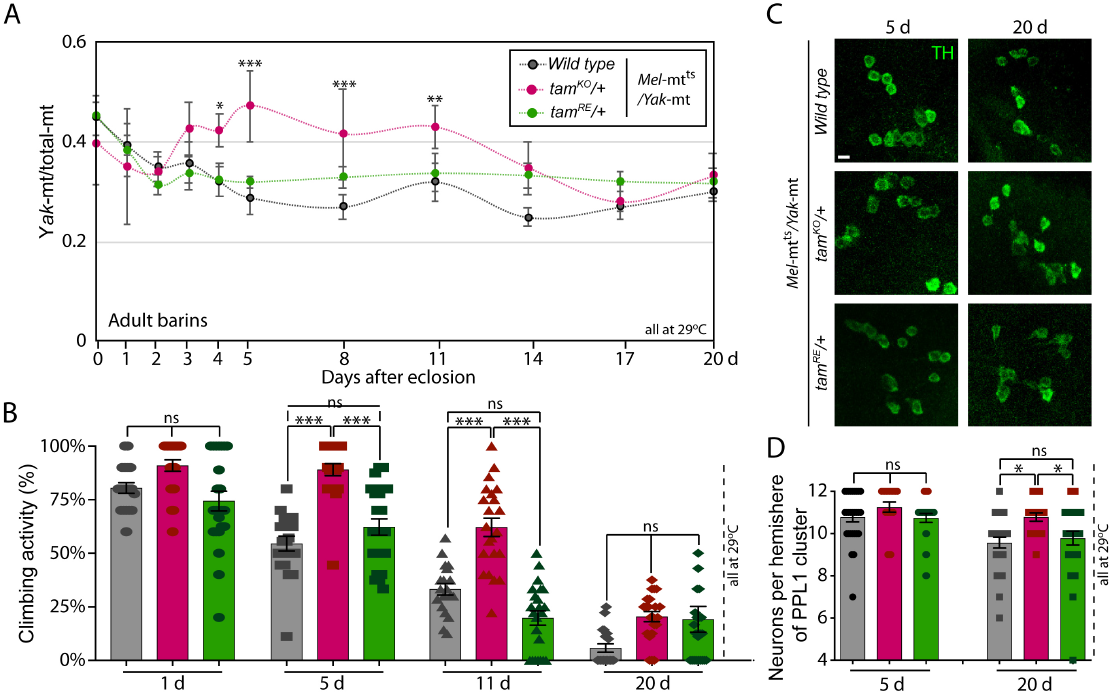
Genetically induced improvements in mitochondrial genome quality and vigor during aging. **(*A*)** A time course during adulthood shows the consequence of removing one of the two alleles of the gene encoding the mitochondrial DNA polymerase (*tam*^*KO*^*/+*) on *Yak-*mt/total-mt (red) compared to two controls (black is wild-type, and green is a revertant *tam* allele). Flies were held at 29°C and sampled at the indicated times. **(*B*)** Removing one allele of mitochondrial DNA polymerase gene suppresses the age-associated decline in climbing activity in heteroplasmic flies. *tam*^*KO*^*/+* in red, wild-type grey and revertant green. **(*C*)** Reducing dose of *tam* in heteroplasmic fly sustains the number of DA neurons (marked by anti-Tyrosine hydroxylase) in the PPL1 clusters of adult brains 20 days after eclosion. **(*D*)** Quantification of DA neuron-number in PPL1 clusters.

### Kinetin treatment enhances purifying selection during aging in flies and mice

Encouraged by findings that genetic manipulations altered the balance of mitochondrial genomes, we wondered whether pharmaceutical activation of quality control might reverse mutational accumulation in aging animals. Kinetin is a modified form of adenine and a cytokine in plants (29). Kinetin and its ribosyl derivative have been shown to activate PINK1 in mammalian cells (30, 31). We tested the influence of kinetin during aging of adult flies. We fed 3-day old adults vehicle control (DMSO), the natural purine (adenine, 100µM) or kinetin (100µM), and followed the ratio of mitochondrial genomes (Fig. 4A). While *Yak*-mt/total-mt was relatively stable in controls, the ratio rose progressively in the kinetin fed flies. The effect of kinetin was dose-dependent, and neutralized by adenine competitor (*SI Appendix*, Fig. S6). We conclude that kinetin treatment promotes purifying selection in our heteroplasmic line during aging.

**Figure 4.**
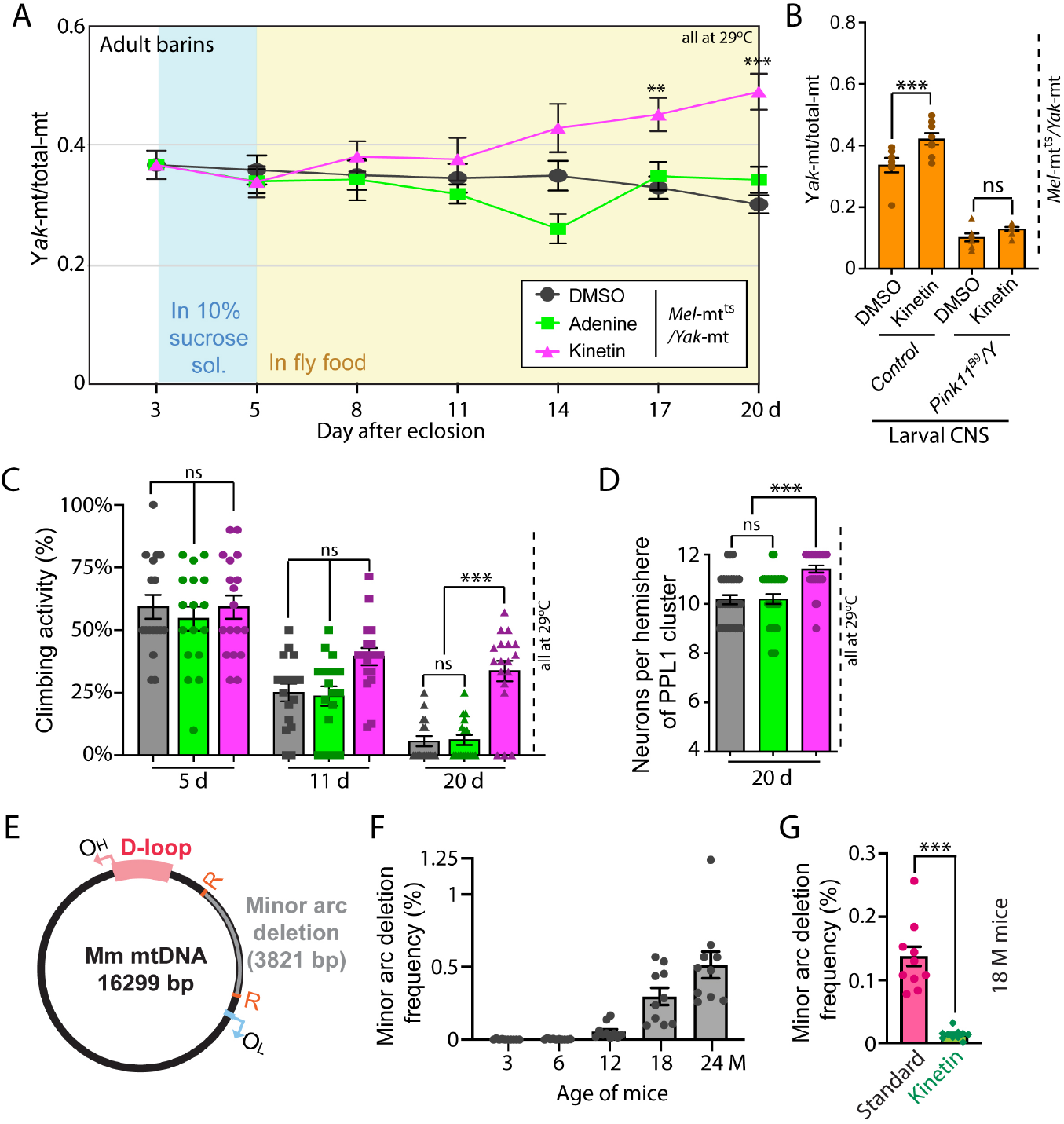
Pharmacological enhancement of PINK1 improves mitochondrial genome quality and vigor. **(*A*)** Kinetin treatment increases *Yak-*mt/total-mt in adulthood. Wild type adult male heteroplasmic flies at 29°C were fed (Methods) with solvent control (DMSO), 100 μM kinetin or 100 μM adenine from day 3 and *Yak-* mt/total-mt was measured in brains at the times indicated. **(*B*)** Kinetin induced increase in *Yak-*mt/total-mt required *Pink1. Yak-*mt/total-mt increase in the CNS of control late 3^rd^ instar larvae (29°C) that were fed kinetin, but not larvae lacking *Pink1, Pink1*^*B9*^*/Y*. **(*C*)** Kinetin sustains the climbing activity in heteroplasmic flies with age. Heteroplasmic males at 29°C were fed with control (DMSO, grey), 100 μM adenine (green) or 100 μM kinetin (red) from day 3, and tested for climbing activity at the indicated times. **(*D*)** Kinetin treatment prevents neuronal loss in PPL1 clusters of heteroplasmic males. Quantification of neuron number of PPL1 clusters in each hemisphere in heteroplasmic male adult flies at 29°C fed with DMSO, 100 μM adenine or 100 μM kinetin from day 3 and assayed at day 20. **(*E*)** Illustration of deletion (grey) located between two 9-bp repeat sequences (orange “R”) within minor arc. **(*F*)** Age dependency of deletion frequency in liver of C57BL6/J mice. Two samples from each of 5 mice were measured by ddPCR for each age group. **(*G*)** Kinetin effect on minor arc deletion levels in aged mouse liver. 16.5 months old mice were fed chow supplemented with kinetin for 6 weeks. This treatment reduced minor arc deletion levels 10-fold compare to control chow-feed mice of the same age.

Since *Pink1*-mutant heteroplasmic flies do not survive into later adulthood, to test whether the kinetin action depends on PINK1 function, we examined heteroplasmic larvae. Kinetin (50µM) feeding increased *Yak*-mt/total-mt in the CNS of control heteroplasmic larvae at 29°C. This action of kinetin depended on *Pink1* consistent with the expectation that kinetin acts by stimulating PINK1 (Fig. 4B). Furthermore, while kinetin increased *Yak*-mt/total-mt in control heteroplasmic larvae at 29°C, it had no effect on larvae at 22°C (*SI Appendix*, Fig. S6c), suggesting that the effect requires the functional distinction between the genomes as expected if kinetin acts by stimulating quality control.

We then tested whether increased *Yak*-mt/total-mt in older kinetin treated flies is associated with improved well-being. Kinetin-feeding sustained the climbing activity (Fig. 4C), and maintained the DA neurons in PPL1 in aged heteroplasmic flies (Fig. 4D). Thus, kinetin treatment promotes purifying selection through a PINK1 dependent mechanism and mitigates the decline of vigor of heteroplasmic flies.

As in the fly model, competition between mutant and wild-type mitochondrial genomes ought to influence accumulation of mutations in mammals. If failing capabilities of purifying selection in aged mammals contribute to age-associated accumulation of mutations as in flies, perhaps kinetin treatment would suppress or reverse age-associated accumulation of a mitochondrial mutation in mice.

We used a sensitive digital PCR assay to measure age-associated accumulation of a naturally emerging deletion in the minor arc of mtDNA in liver of WT C57BL6/J mice (Fig. 4F). Note that because mutations are rare events first emergence of the deletion will vary from mouse to mouse leading to some quantitative variation. Nonetheless, mutational abundance rises dramatically in older mice, with near exponential response to age. We fed control or kinetin containing food to 16.5-month-old mice for 6 weeks and assessed the levels of the minor arc deletion in liver. Kinetin feeding reduced the level of the minor arc deletion well below similarly aged mice fed control food (Fig. 4H). Moreover, kinetin reduced the mutation load below the level characteristic of the age at onset of kinetin feeding (*SI Appendix*, Fig. S7). This shows that a PINK1 activator reverses the age-associated increase of this mutant allele.

## Discussion

Random segregation of mitochondrial genomes can lead to stochastic variation in the relative abundance of coresident genomes within the cells of an individual (6, 32), but selective forces often bias the outcome to drive directional changes (6, 19, 20, 22, 32–35). In the female germline of *Drosophila*, quality control promotes biogenesis and proliferation of those mitochondria that maintain a useful membrane potential (18, 19, 36). This puts mitochondria compromised by deleterious mutations at a proliferative disadvantage resulting in their elimination within a few generations. It has, however, been less clear how selective forces and quality control mechanisms influences the abundance of mutant genomes in the soma. The accumulation of mitochondrial mutations is important in the progression of heteroplasmic mitochondrial disease and impacts normal aging.

A genetic model for heteroplasmy in *Drosophila* provided a means of assessing selective forces that influence the abundance of a deleterious mutation. While the overall abundance of the genome with the deleterious mutation is influenced by a variety of selective forces, by using a temperature sensitive mutation and assessing the results at two different temperatures, we can isolate the influence of the change in function of the *mt:CoI* gene product. We take the difference in the abundance at the two temperatures as a measure of the impact of purifying selection. By introducing mutations altering quality control pathways, we can assess the role of the affected pathway in purifying selection. This allowed us to show that purifying selection operates in somatic tissues during development and growth (Fig. 1), and that a PINK1 dependent pathway contributes importantly to this purifying selection (Fig. 2A). Sensitivity to the reduction in the dose of *larp* and enhancement by reduction in the dose of *tam* suggest that the mechanism operating in the soma is similar to that described for purifying selection during oogenesis (18), but there may be additional mechanisms contributing.

Selection can only influence the relative proportion of competing mitochondrial genomes if there is replication and/or turnover of the genomes. Thus, stability might contribute to the absence of observed purifying selection in the adult. However, this cannot explain periods of dynamic change in *Yak*-mt/total-mt. For example, the *Yak*-mt/total-mt ratio in the gut falls during adulthood, but falls the same amount at both temperatures (Fig. 1F). Additionally, we characterized a very rapid decline in *Yak*-mt/total-mt in the young adult following eclosion (Fig. 1D,E). Because this decline erases the benefits of earlier purifying selection, we interpret the change as the dynamic resetting of *Yak*-mt/total-mt ratio after a major reduction in the effectiveness of purifying selection.

We measure the impact of purifying selection rather than the process itself. Consequently, a rise in a counteracting selection, rather than a loss in purifying selection might be responsible for the reduction in the impact of purifying selection. The unexpected prevalence of deleterious mutations in adult mutator flies led to the suggestion that such mutations might be positively selected (23). Although, we do not know how such a “destructive” selection would occur, if it exists, it would act in opposition to purifying selection and so reduce its impact. Accordingly, it is possible that a rise in destructive selection accounts for our recorded changes in adulthood.

Perhaps our most impactful finding is that purifying selection can be at least somewhat restored in later adulthood either genetically or pharmacologically (Fig. 3,4). In addition to improving mitochondrial genome quality these treatments enhanced the climbing ability of aged heteroplasmic flies and suppressed loss of dopaminergic neurons (Fig. 3,4). This ability to reverse negative effects of heteroplasmy on the aging of flies suggests that such treatment might have the potential to reverse detrimental consequences of heteroplasmic mitochondrial diseases and perhaps suppress age-associated accumulation of mitochondrial mutations. Such possibilities are further bolstered by our finding that kinetin, the pharmacologic agent promoting purifying selection in adult flies, also works in mice to suppress age-associated increase in a detrimental mtDNA mutation (Fig. 4).

Our findings raise questions of how the quality control mechanisms decline with age. A previous study, using a different model that assesses the levels of a deletion of mtDNA that was induced in adult *Drosophila* flight muscle also found negligible levels of purifying selection (22). This study went on to demonstrated that a number of genetic alterations enhanced purifying selection. This led to the conclusion that “wild-type levels of key gene products are not set to maximize mtDNA quality control”. Specifically, this work implied that such processes as mitochondrial fusion and fission as well as the ability of the mitochondrial ATP synthase to run in reverse are not optimized for quality control in the adult. Together with our findings, this report suggests a number of different pathways that are candidates for pharmacological modification that will benefit mtDNA quality during aging. While the findings also suggest that many different pathways might limit quality control in the adult, it does not explain why function is not optimized.

While it is puzzling why there is a lapse of purifying selection in the adult, we suggest that it may be the fallout of the shifting priorities of evolution at different stages of life. Growth and development of metazoans occurs in protected stable environments where organismal performance is not a factor, but maintenance of genome quality is key to the success of the future individual and to production of future generations. Under these circumstances evolution succeeds by prioritizing purifying selection. In contrast, the survival and reproductive potential of the adult depends on performance in a challenging and varied environment. At this life stage, evolution would select for elite performance, and might optimize the energy-producing capacity of mitochondria by increasing the DNA content of weak mitochondria, even at the expense of triggering amplification of less functional genomes.

Regardless of the underlying reasons, we show a decline in the impact of quality control in the adult, and that a decline in vigor parallels reductions in the average quality of mitochondrial genomes during aging. These observations, together with a substantial literature (6, 10–12, 18–22, 25, 32–35), suggest that the mitochondrial “genotype” of a metazoan is plastic and molded by selective forces acting within the organism. Importantly, we show that mitochondrial genome quality can be genetically and pharmacologically modified during an organism’s lifetime to benefit well-being (Fig. 3, 4). These findings bode well for development of therapies that modulate selective forces to enhance mitochondrial quality to benefit healthspan and lessen the impact of mitochondrial disease.

## Materials and Methods

### Fly stocks

The heteroplasmic fly (*Mel-*mt^ts^*/Yak-*mt) containing both a *D. melanogaster* mt genome with mutant two alleles (*mt:ND2*^*del1*^ + *mt:CoI*^*T300I*^) and a *D. yakuba* mitochondrial genome were previously described (25). We are using a derived stock of the original heteroplasmic line (25) that carries a higher proportion of *Yak-mt. Mel-*mt^ts^*/Yak-*mt females were backcrossed to males of a laboratory version of Canton S for 10 generations to homogenize the nuclear background (the lab stock was itself created by ingression of a Canton S background into *w1118*). The *tam*^*RE*^, and *tam*^*KO*^ alleles are congenic transgenic flies with a restored wild-type allele of *tam* (control) and a knockout allele, respectively (gifts from Dr. Wredenberg (37)). *larp*^*mtr-null*^ originated from Dr. Glover and was a gift from Dr. Xu(17). *tam*^*3*^, *tam*^*4*^, *PINK1*^*RV*^, *PINK1*^*B9*^ and *Dp(1:3)* ^*DC026*^ were obtained from the Bloomington Drosophila Stock Center. The stocks were cultured at 22°C and 29°C on standard fly medium.

### Mice

Male C57BL/6J mice were obtained from The Jackson Laboratory, housed in a specific pathogen-free facility with a standard 12 h light/dark cycle at the University of California, San Francisco and given food and water ad libitum. Experiments were conducted in accordance with institutional guidelines approved by the University of California, San Francisco Institutional Animal Care and Use Committee.

### DNA isolation

Total DNA of adult flies was extracted as follows. Three adult flies were pooled and mechanically homogenized with a plastic pestle in 105 μl homogenization buffer (100 mM Tris-HCl [pH 8.8], 0.5 mM EDTA, 1% SDS). The homogenate was incubated at 65° for 30 min, followed by addition of 15 μl potassium acetate (8 M) and incubation on ice (30 min) to precipitate protein and SDS. Subsequently, the homogenate was centrifuged at 13,000 rpm for 15 min at 4°. DNA was recovered from the supernatant by adding 0.5 volumes of isopropanol and centrifuging at 20,000 g for 5 min at room temperature. The resultant pellet was washed with 70% ethanol and suspended in 20 μl of ddH_2_O. DNA extracted from tissue samples as follows. A single dissected fly tissue (brain, wing disc or gut) was placed on a clean cover slide, mixed with 10 μl of lysis buffer (10 mM Tris-HCl [pH 8.8], 1 mM EDTA, 25 mM NaCl, 1% SDS and 8U/ml Proteinase K, NEB) for 3 min at room temperature. The tissue/buffer mixture was transferred to a 0.2ml tube and incubated at 37° for 30 min and heat inactivated by 95° for 5 min.

Mice were anesthetized by intraperitoneal injection of 0.3cc 17.5 mg/ml Ketamine and 10 mg/ml Xylazine mixture in saline and perfused with 15-20 ml sterile PBS through heart left ventricle using the surflo wing infusion set (Terumo Corporation, SV-21BLK) attached to 30 ml syringe. After perfusion livers were collected, immediately frozen on dry ice and stored at –80 °C until DNA isolation. Two small liver fragments (~1mm^3^) per animal were analyzed. Fragments were lysed in 90 μl of lysis solution (50 mM Tris-HCl pH 8.0, 10 mM EDTA pH 8.0, 0.5% SDS, 2 mg/ml proteinase K (Promega, v3021)) at 37 °C overnight followed by addition of 10 μl of 3M potassium acetate and incubation at room temperature for 1h. Next, lysates were centrifuged at 21000 g for 10 min and supernatants were transferred to silica membrane spin columns (Macherey-Nagel) and centrifuged at 11000 g for 1 min. Flow-throughs were heat inactivated at 70 °C for 30 min and cleaned up using home-made magnetic beads (1:1 ratio, (38)). After clean-up DNA was eluted in 0.1x TE buffer and stored at 4 °C.

### qPCR analyses

For all qPCR assays, SYBR^®^ Select Master Mix (Applied Biosystems 4470908) was used in 20 μl reactions with 400 nM of each primer. To measure the total mtDNA copy number of heteroplasmic flies, qPCR of a 128 bp *mt:IrRNA* region present in both mtDNA genotypes were performed (primer AACCAACCTGGATTACACCG and TGCGACCTCGATGTTGGATT) and normalized to nuclear genome copy number of the *αTub84D* gene (primer ATGCGCGAATGTATCTCTATCC and AGGTGTTAAACGAGTCATCACC). To measure copy number of *D. yakuba* genomes, qPCR of the 71-bp *D. yakuba mt:COXI* region was performed (primer AGTTGAAAACGGAGCTGGTACA and CTCCACCATGAGCGATACCTG). The efficiency and specificity of *D. yakuba mt:COXI* and total *mt:IrRNA* primer sets in qPCR reaction was normalized each time by analyzing the ratio between the two primer sets in DNA samples from a *D. melanogaster* wild type stock of Canton S. The background for the *Yak-*mt specific reaction was also assessed in every qPCR experiment and was always less than 0.005% of total-mt in DNA samples from Canton S. qPCR was performed with the following reaction conditions: 95° for 3 min, followed by 40 cycles at 95° for 5 s and 62.5° for 15 s. For each 20μl qPCR reaction, 0.375% of an adult fly’s, or 0.667% of a tissue’s total genomic DNA was used as template. The percentage of *D. yakuba* mtDNA was calculated by dividing *D. yakuba* mtDNA copy number by the total mtDNA copy number. The Ct values used ranged from 18 to 30.

### ddPCR analyses

The ddPCR analysis was performed on a QX100 system (BioRad). Two assays designed to detect minor arc deletion (primer LD_F CCAGAAGATTTCATGACC, primer LD_R CTTCTAGGTAATTAGTTGGG, probe LD AACTAATCCTAGCCCTAGCC) and common sequence for both deleted and non-deleted mitochondrial genomes (primer LD_common_F TTAGGATATACTAGTCCGC, primer LD_common_R GAAGTCTTACAGTCCTTATC, probe LD_common AGCCTTCAAAGCCCTAAGA) were used simultaneously. Template DNA concentration was adjusted to be below 3500 mitochondrial genome copies per μl of ddPCR reaction mixture. The reaction mixture (total volume 22 μl) contained 11 μl of 2× ddPCR Super Mix for Probes (BioRad, 1863024), 1.1 μl of 4 primers mixture 18 μM each, 1.1 μl of 5 μM probe LD_common labeled with HEX (IDT), 1.1 μl of 5 μM probe LD labeled with FAM (IDT), 0.5 μl of Alu1 restriction enzyme (NEB, R0137L), 2.2 μl of ddH_2_O and 5 μl of template DNA. 20 μl of the reaction mixture and 70 μl of oil (BioRad, 1863005) were loaded on a DG8 cartridge (BioRad, 1864007) for droplet generation on QX100 Droplet Generator (BioRad). 40 μl of droplet emulsion were transferred to 96-well plate (Bio-Rad, 12001925) and sealed with a pierceable foil (BioRad, 1814040) using PX1 PCR plate sealer (BioRad). The optimized PCR thermal cycling was conducted on a conventional PCR machine (BioRad, C1000 Touch) using the following conditions: 10 min activation period at 95 °C, followed by 40 cycles of denaturation at 94 °C for 30 s, ramp rate 2 °C, and combined annealing-extension at 54°C for 1 min, ramp rate 2 °C, and 1 cycle of 98 °C for 10 min. After thermocycling samples were analyses on QX100 Droplet Reader (BioRad). Results were analyzed with the QuantaSoft Analysis Pro v.1.0.596 software (BioRad).

### Behavior Assay

Climbing ability was defined as the ability of the adult fly to climb 5 cm within 10 s. If the fly could accomplish the task (≤ 10 s), it was given a score of 1; otherwise, it was given a score of 0 (≥ 10 s). Ten flies collected from an independent cross were grouped into a single task. 15~25 tasks (150~250 flies from 15-25 independent crosses) were tested of each genotype.

A number of independent crosses, 15, 22 or 25 were carried out to produce flies of each genotype, wild-type, tam^KO^/+, or tam^RE^/+, respectively. Ten male flies were collected from each cross to form independent cohorts (62). Each cohort was challenged in a climbing test done at day 1, 5, 11 20. In each test climbing ability was defined as the ability of the adult fly to climb 5 cm within 10 s. If the fly could accomplish the task (≤ 10 s), it was given a score of 1; otherwise, it was given a score of 0 (≥ 10 s). All surviving flies were scored, and the % success of the surviving flies was tabulated and plotted. In total, 248 independent tests were conducted.

## Check numbers with Pei-I

### Immunocytochemistry and Confocal Microscopy

Adult brains were dissected in PBT (0.3% Tween 20 in PBS), and incubated with fixation solution (4% formaldehyde in PBT) for 20 min, followed by blocking for 1 h with 1% BSA in PBT. Samples were immunostained with rabbit anti-Tyrosine hydroxylase (AB152; EMD Millipore Corporation) at 1:200, and Alexa 488-conjugated anti-rat (ab150165; AbCam. Samples were imaged with a 20X dipping 2mm WD objective on a Zeiss Confocal Laser-Scanning Microscope 780 with identical imaging parameters among different genotypes in a blind fashion. Images were processed with Photoshop CS4 using only linear adjustment of contrast.

### Kinetin Treatment in fly

For larval treatment, 0-6 h eggs were collected and grown on regular fly food (10 ml) containing control DMSO or 50 μM Kinetin. Six h after egg-collection, 10 μl of DMSO or 5 mM kinetin (in DMSO) was mixed with 190 μl ddH_2_O and applied on the top of the fly food for control and experimental larvae, respectively. Tissues were dissected from late 3^rd^ instar larvae. For adult treatment, 3-d old male adults were collected and kept in vials with a cotton ball moistened with 2 ml of 10% sucrose water solution with DMSO, 100 μM kinetin, 100 μM Adenine, or 100 μM kinetin & 100 μM adenine for 48 h. Then, the flies were transferred to the vials as described below and passaged to similar fresh vials every 2 days until dissection. Recipient vials with 10 ml of regular fly food were prepared fresh by adding 100 μM kinetin, 100 μM adenine, or 100 μM kinetin & 100 μM Adenine in 10 μl of DMSO, 10 μM kinetin, 10 μM adenine, or 10 μM kinetin & 10 μmM Adenine in 190 μl ddH_2_O, respectively, on the surface of the fly food, and air-dried.

### Kinetin chow

Kinetin (Sigma) was delivered orally to mice in their chow following published reports (39). Rodent chow (Purina 5053) was formulated by Research Diets (New Brunswick, NJ) to contain 3.50 g kinetin per kg chow for mice. These amounts of kinetin were well tolerated during trials according to a previous report (39). Chow was stored at −80°C. Fresh chow was provided at least every four days. The delivery of chow was not blinded with respect to drug treatment. The 16.5-month-old mice were fed with kinetin or control chow for 6 weeks before sample collections at 18 months of age. Body weights and chow intake were monitored at least twice weekly.

## Acknowledgments

We thank Chun-Yi Cho of the O’Farrell laboratory for insightful discussions and suggestions on this manuscript. This work was supported by a National Institutes of Health grant R35GM136324 and by a Larry L. Hillblom foundation grant 2019-A-011-NET to P.H.O’F.

## Supplementary Information for

**Fig. S1.**
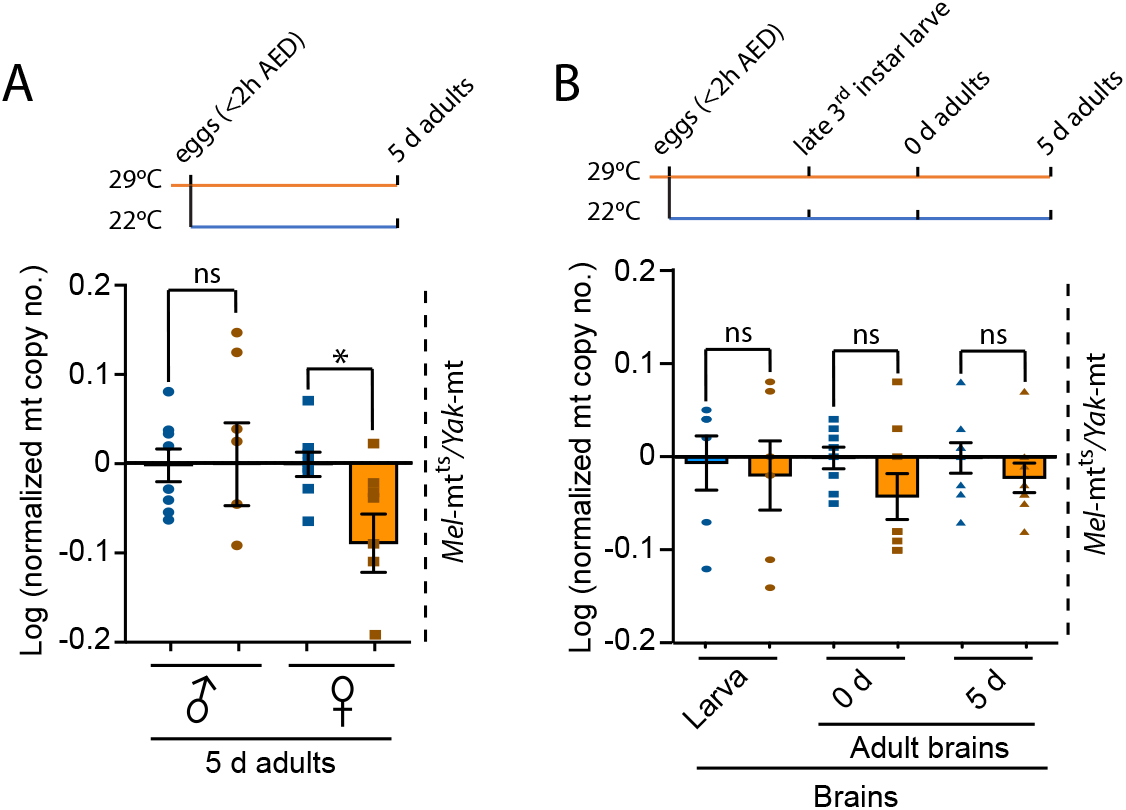
Activating quality control by temperature inactivation of *mt:CoI*^*ts*^ does not alter total mt copy number. The log of a ratio of qPCR measurements (here and below) represent up and down changes in mtDNA copy number symmetrically around zero. The left sample acts as the control (normalizing) value. **(*A*)** In contrast to the influence of temperature on mt genome quality (Fig. 1), an increase in the temperature of development from 22 to 29°C did not alter the total copy number in males. The decrease in copy number in females at 29°C is likely due to reduced ovarian production of oocytes at the high temperature. Three adults were grouped as one biological sample and 8 biological repeats were measured for each condition. **(*B*)** Temperature was not associated with significant change in mt copy number in the brain. In the three pairs of bars, the 29°C data is normalized to the 22°C data for the same stage to show an absence of temperature effect. For each time and condition in **(*B*)** each data point is a single whole brain from one individual and 8 data points were collected at each time and stage. * = *p* < 0.05 by one-way ANOVA/Tukey’s multiple comparison test.

**Fig. S2.**
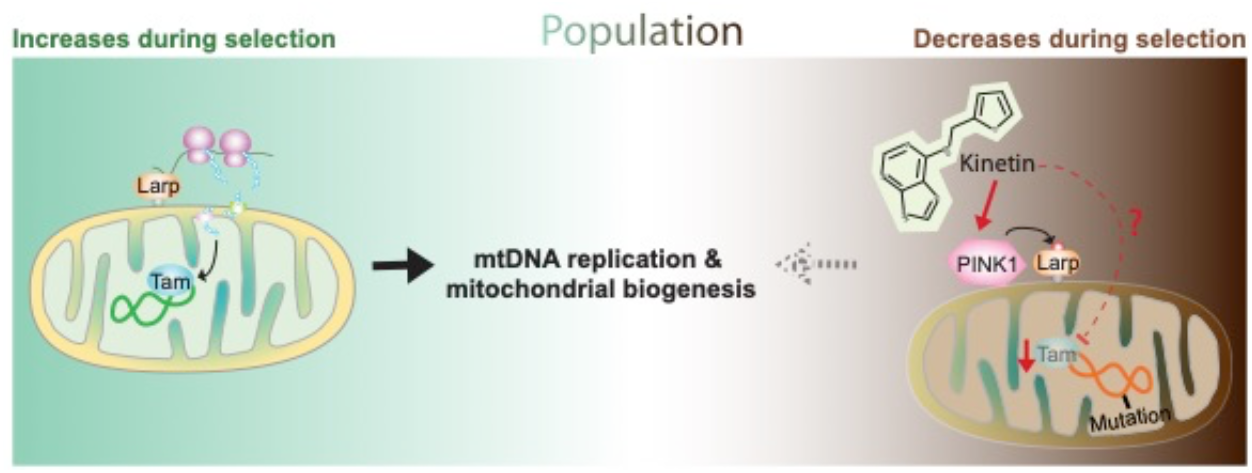
Model for Kinetin/PINK1 regulation through Tamas in selective amplification of mitochondrial genomes in somatic tissues. Mitochondrial biogenesis is driven largely by translation of nuclear encoded mRNAs whose protein products are transported into mitochondria. The targeting of relevant mRNAs to mitochondria and local translation facilitate multiplication of mitochondria. Larp-based recruitment of appropriate mRNAs to the mitochondria underlies this local translation. The mRNA for the mitochondrial DNA polymerase, POLG (Tam in *Drosophila*), is among the mRNAs recruited to mitochondria. As described by Zhang et al., 2019, the system discriminates against unhealthy mitochondria, which accumulate the serine/threonine kinase PINK1 on the surface where it phosphorylates Larp, inhibiting its ability to recruit mRNAs for local translation. The result is selective mitochondrial biogenesis that enhances proliferation of healthy mitochondria giving them a selective advantage. While the system impacts the local translation of many mitochondrial proteins, in situations in which Tam is limiting, its selective translation might directly promote selective replication of genomes associated with healthy mitochondria (dashed red line). If other mitochondrial proteins are limiting, coupling of mtDNA replication to biogenesis ought to indirectly produce the same selective pressure. In a dynamic population with turnover, this selective biogenesis will progressively replace mutant mitochondria with healthy mitochondria, a form of intracellular purifying selection. However, when this purifying selection weakens with age, proliferation of defective mitochondrial genomes will no longer be constrained. By treating with Kinetin to re-activate PINK1 in elderly animals, we enhance quality control to restore purifying selection, improve the mean quality of the population of mitochondrial genomes, and improve vigor.

**Fig. S3.**
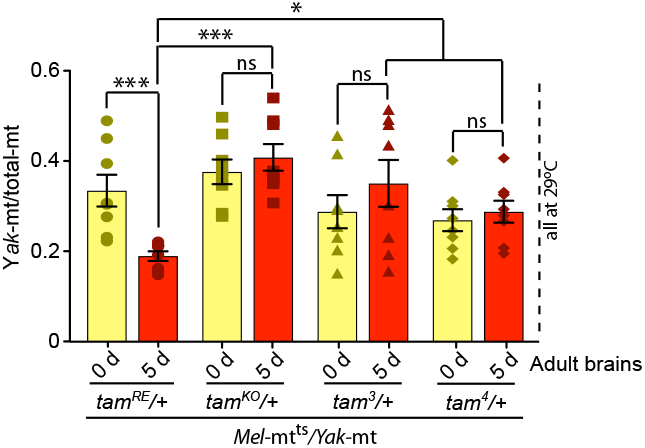
Different alleles of *tam* all reduce the age-associated drop in *Yak*-mit/total-mt. Different mutant alleles of *tam* (*tam*^*KO*^, *tam*^*3*^ and *tam*^*4*^) all cause persistence of mitochondrial genome quality during early adulthood when heterozygous. *tam*^*RE*^ is a revertant allele congenic with *tam*^*KO*^ that is used as a control. Bars show the average ratio of *Yak-*mt to total-mt in brains dissected from newly eclosed (yellow) or 5-day old (red) adults of the stated genotypes that were raised at 29°C. For each stage, time and genotype, 8 brains were collected from individuals for mtDNA analyses. * = *p* < 0.05, ** = *p* < 0.01, and * = *p* < 0.001 by one-way ANOVA/Tukey’s multiple comparison test.

**Fig. S4.**
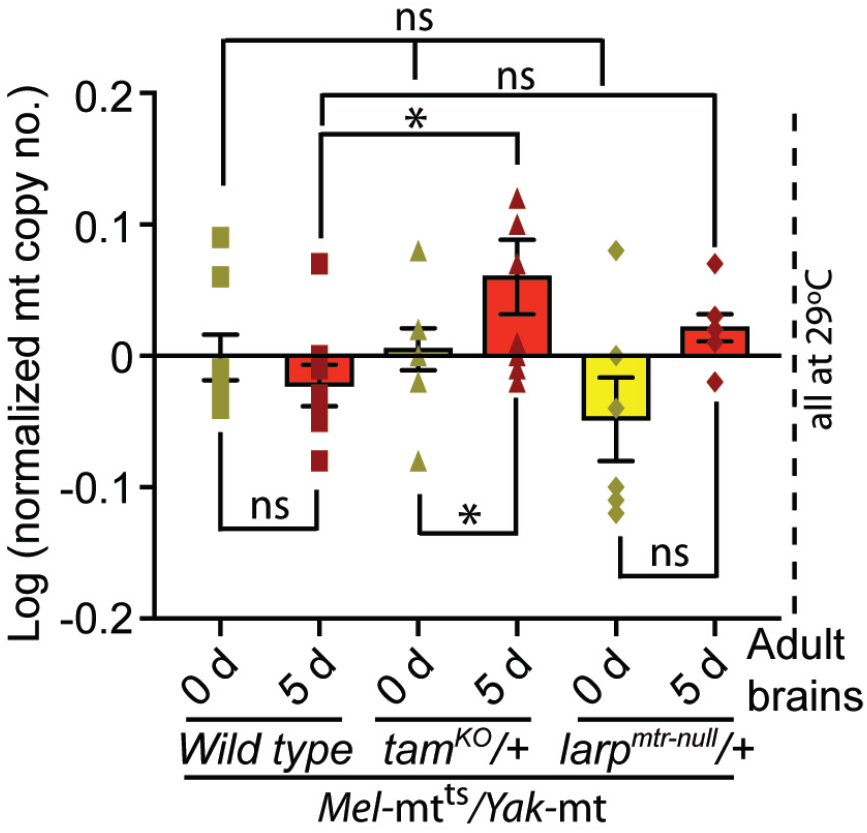
Dosage of quality control genes have minimal effects on mtDNA copy number in young adults. We measured total mt copy number at eclosion (0 d) and at 5 days of adulthood (5 d) in heteroplasmic strains with a nuclear genome that was wild type or heterozygous for either *tam* or *larp*, as indicated. The flies were maintained at 29°C. Compared to newly eclosed wild type, small shifts were observed in the copy number of the heterozygous mutant lines, but only the slightly increased level in *tam*^*KO*^/+ scored as modestly significant. For each time and genotype, we tested DNA from 8 whole brains that were collected from different individuals. * = *p* < 0.05 by one-way ANOVA/Tukey’s multiple comparison test.

**Fig. S5.**
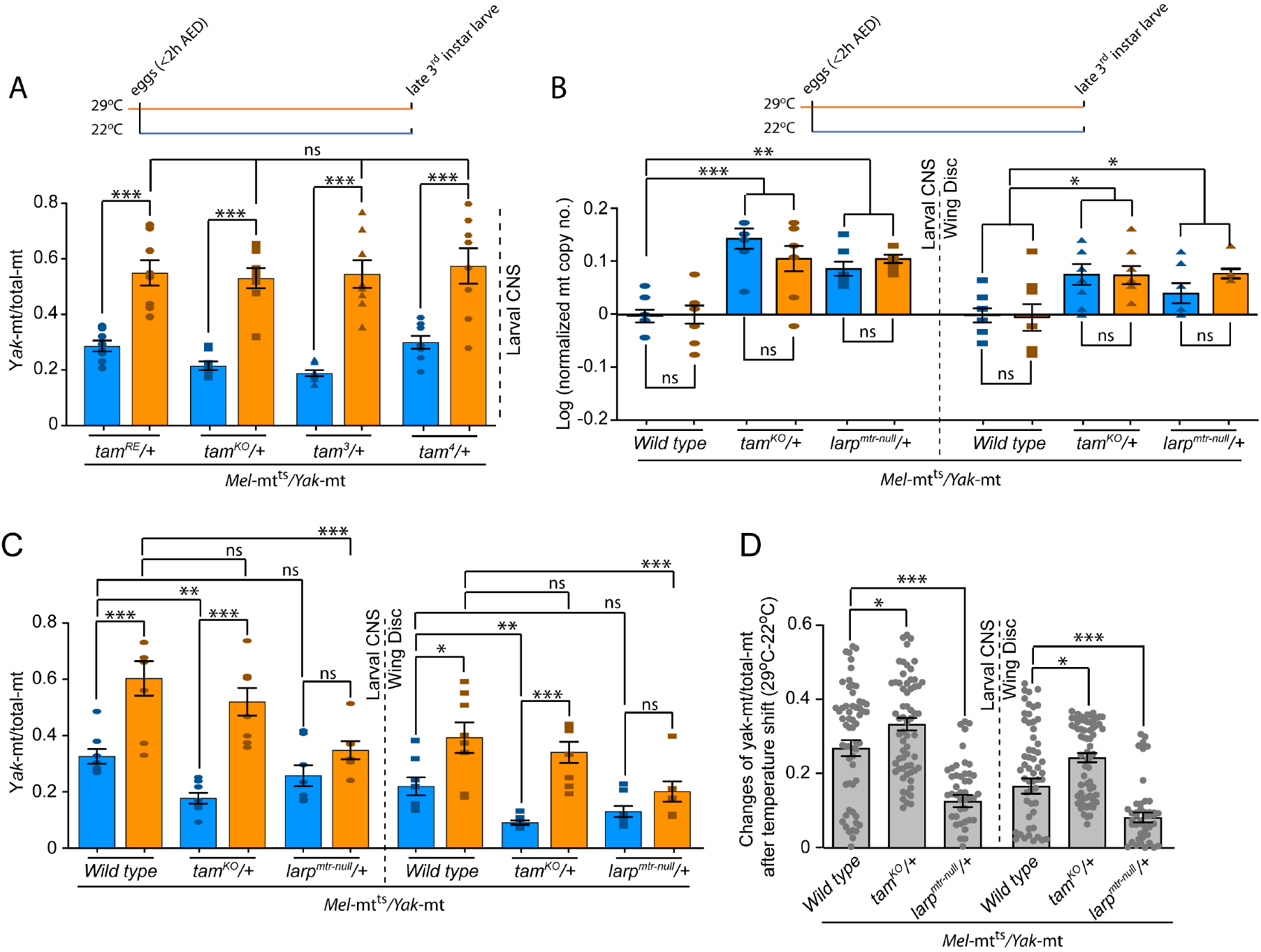
Nuclear genotype has multiple influences on larval heteroplasmy. **(*A*)** Unlike the results in the 5-d adult, reduction of the dose of functional *tam* (3 different alleles) did not alter *Yak*-mt/total-mt at 29°C at the larval stage (amber). However, some variation in the ratio occurred at 22°C that might be due to effects of *tam* heterozygosity (congenic control-*tamRE* vs *tam*^*KO*^) as well as genetic background (*tam*^*3*^ vs *tam*^*4*^). **(*B*)** Heterozygosity for *tam* or *larp* increased the copy number of total mtDNA irrespective of temperature. Shown for two larval tissues, wing disc and CNS from heteroplasmic stains (*Mel-mt*^*ts*^*/Yak-mt*) with indicated nuclear genotypes raised at 22°C (blue) or at 29°C (amber) and normalized to wild-type 22°C. This suggests that the dose of these genes have a mild effect on copy-number control that is independent of the discordance in the functionality of the two genomes. **(*C*)** Nuclear genotypes have a complex influence on the ratio of *Yak*-mt/total-mt in larval tissue. The ratio of mt genomes was measured in two larval tissues (CNS and wing disc), at two temperatures (22°C and 29°C) and for the three nuclear backgrounds, wild type and both heterozygous mutants (*tam*^*KO*^/+ and *larp*^*mtr-null*^/+). Higher temperature increased the *Yak*-mt/total mt ratio (improved mt quality) in all nuclear genotypes but the increase in the *larp* heterozygote fell below significance, and heterozygosity for *tam* decreased the *Yak*-mt/total-mt at 22°C. **(*D*)** The two genes, *tam* and *larp*, have opposite but quantitatively small effects on quality control in these larval tissues. The influence of nuclear genotype on quality control is not easily seen in panel **(*C*)** because of the complicating influence of the nuclear genotype on the ratio of the two mitochondrial genomes at 22°C where both genomes are functional. To isolate the effect on quality control, we plotted the difference between 22°C and 29°C for each genotype, plotting the level at 29°C minus the level at 22°C. The results show that heterozygosity for *tam* modestly enhanced quality control, while heterozygosity for *larp* substantially compromised it, behaviors consistent with the effects of these mutations seen at the adult stage. While **(*C*)** plots 8 data points, in **(*D*)**, each sample (8) at one temperature was compared to every sample (8) measured at the other temperature to give 64 values that are plotted. * = *p* < 0.05, ** = *p* < 0.01, and * = *p* < 0.001 by one-way ANOVA/Tukey’s multiple comparison test.

**Fig. S6.**
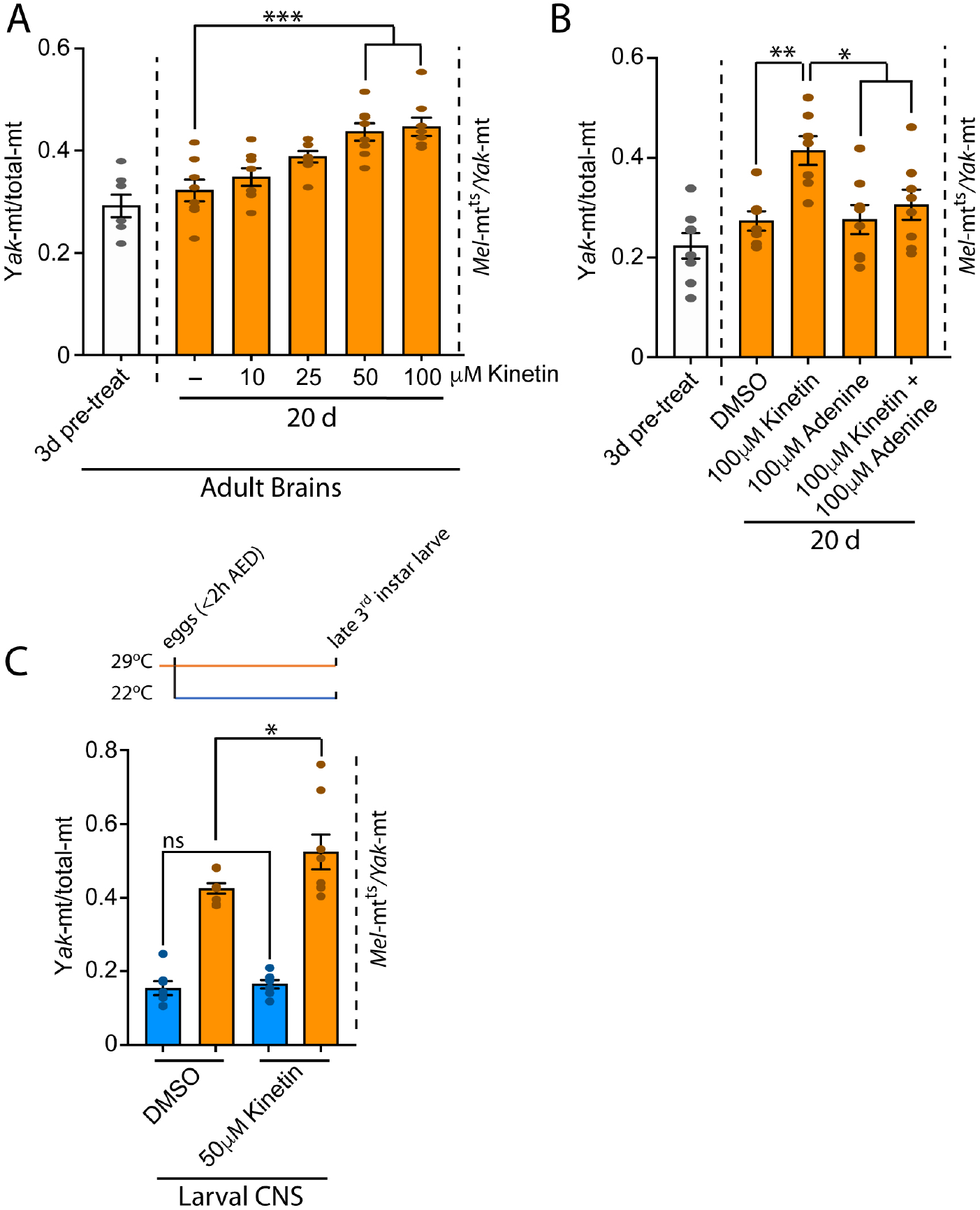
Kinetin acts as a dose-dependent enhancer of mt genome quality control. **(*A*)** Increasing doses of kinetin fed to heteroplasmic flies from 3 days post-eclosion to 20 days improved mt genome quality. Brains were dissected from adult heteroplasmic flies (*Mel-mt*^*ts*^*/Yak-mt*|*wild type*) either at day 3 (pre-treatment control) or day 20 after 17 additional days at 29°C with treatment at different concentrations of kinetin (0-100 μM, as indicated). These brains were then tested for the ratio of *Yak*-mt to total-mt. **(*B*)** Adenine suppresses the effect of the adenosine analog, kinetin. Brains dissected from adult heteroplasmic flies either at day 3 (pre-treatment control), or at day 20 after 17 additional days at 29°C with treatment with control DMSO, 100 μM kinetin, 100 μM adenine, or 100 μM kinetin+100 μM adenine. These were then were assayed for *Yak-*mt/total-mt. Adding adenine naturalized the effect of kinetin treatment on quality control of mt genomes. Tissue from 8 individuals was analyzed for each condition. **(*C*)** Kinetin action in the larvae. Heteroplasmic larvae (*Mel-mt*^*ts*^*/Yak-mt*|*wild type*) were raise at either 22°C or 29°C in the presence of 50 μM kinetin or solvent control (DMSO). The ratio of *Yak-*mt/total-mt was measured in the CNS of late 3^rd^ larvae. * = *p* < 0.05, ** = *p* < 0.01, and * = *p* < 0.001 by one-way ANOVA/Tukey’s multiple comparison test.

**Fig. S7.**
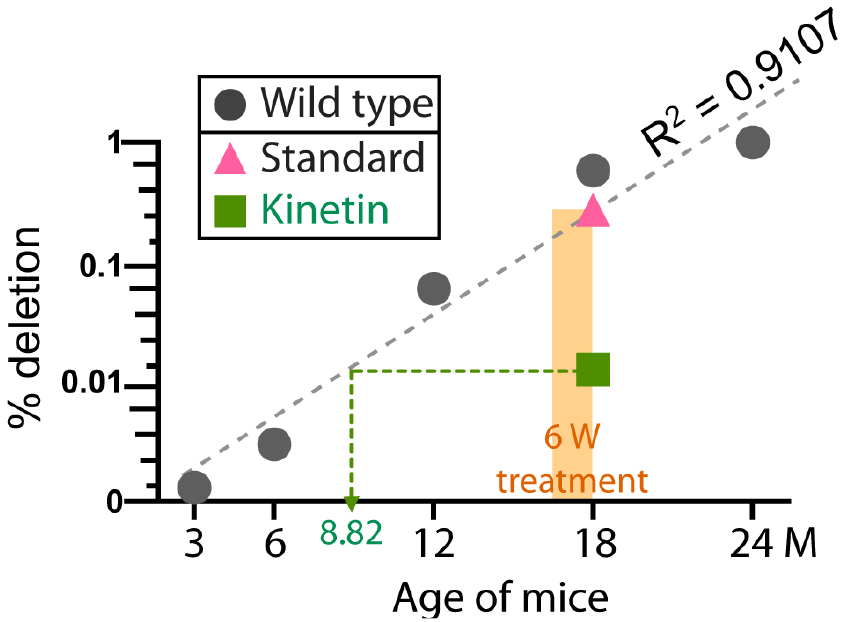
Kinetin feeding reduces mutational load in C57BL6/J mice. Replotting of the data in Fig. 4*F* and 4*G* on a single graph using a logarithmic scale helps visualize the early stages of the age-associated rise in abundance of the minor-arc deletion and the effect of kinetin feeding. In control mice (grey dots), accumulation of the minor arc deletion is roughly exponential over the period of analysis (dashed grey line is a linear regression fit). Kinetin (green ■) feeding for 6 weeks (orange bar) reduces the level of the minor-arc deletion while feeding control food does not (pink Δ). Since the post-treatment level is considerably below the level expected for the onset of kinetin feeding, we suggest that kinetin selects against pre-existing mutations.

## References

1. N. W. Soong, D. R. Hinton, G. Cortopassi, N. Arnheim, Mosaicism for a specific somatic mitochondrial DNA mutation in adult human brain. Nat Genet 2, 318–323 (1992).

2. Y. Michikawa, F. Mazzucchelli, N. Bresolin, G. Scarlato, G. Attardi, Aging-dependent large accumulation of point mutations in the human mtDNA control region for replication. Science 286, 774–779 (1999).

3. D. C. Samuels et al., Recurrent tissue-specific mtDNA mutations are common in humans. PLoS Genet 9, e1003929 (2013).

4. H. L. Baines et al., Similar patterns of clonally expanded somatic mtDNA mutations in the colon of heterozygous mtDNA mutator mice and ageing humans. Mech Ageing Dev 139, 22–30 (2014).

5. A. Trifunovic et al., Premature ageing in mice expressing defective mitochondrial DNA polymerase. Nature 429, 417–423 (2004).

6. J. P. Jenuth, A. C. Peterson, E. A. Shoubridge, Tissue-specific selection for different mtDNA genotypes in heteroplasmic mice. Nat Genet 16, 93–95 (1997).

7. J. B. Stewart, C. Freyer, J. L. Elson, N. G. Larsson, Purifying selection of mtDNA and its implications for understanding evolution and mitochondrial disease. Nat Rev Genet 9, 657–662 (2008).

8. T. Tatsuta, T. Langer, Quality control of mitochondria: protection against neurodegeneration and ageing. EMBO J 27, 306–314 (2008).

9. A. J. Whitworth, L. J. Pallanck, The PINK1/Parkin pathway: a mitochondrial quality control system? J Bioenerg Biomembr 41, 499–503 (2009).

10. C. Vives-Bauza et al., PINK1-dependent recruitment of Parkin to mitochondria in mitophagy. Proc Natl Acad Sci U S A 107, 378–383 (2010).

11. S. Geisler et al., PINK1/Parkin-mediated mitophagy is dependent on VDAC1 and p62/SQSTM1. Nat Cell Biol 12, 119–131 (2010).

12. D. P. Narendra et al., PINK1 is selectively stabilized on impaired mitochondria to activate Parkin. PLoS Biol 8, e1000298 (2010).

13. A. C. Poole et al., The PINK1/Parkin pathway regulates mitochondrial morphology. Proc Natl Acad Sci U S A 105, 1638–1643 (2008).

14. Y. Yang et al., Pink1 regulates mitochondrial dynamics through interaction with the fission/fusion machinery. Proc Natl Acad Sci U S A 105, 7070–7075 (2008).

15. H. Deng, M. W. Dodson, H. Huang, M. Guo, The Parkinson’s disease genes pink1 and parkin promote mitochondrial fission and/or inhibit fusion in Drosophila. Proc Natl Acad Sci U S A 105, 14503–14508 (2008).

16. A. M. Pickrell, R. J. Youle, The roles of PINK1, parkin, and mitochondrial fidelity in Parkinson’s disease. Neuron 85, 257–273 (2015).

17. Y. Zhang, Y. Chen, M. Gucek, H. Xu, The mitochondrial outer membrane protein MDI promotes local protein synthesis and mtDNA replication. EMBO J 35, 1045–1057 (2016).

18. Y. Zhang et al., PINK1 Inhibits Local Protein Synthesis to Limit Transmission of Deleterious Mitochondrial DNA Mutations. Mol Cell 73, 1127–1137 e1125 (2019).

19. H. Ma, H. Xu, P. H. O’Farrell, Transmission of mitochondrial mutations and action of purifying selection in Drosophila melanogaster. Nat Genet 46, 393–397 (2014).

20. J. H. Hill, Z. Chen, H. Xu, Selective propagation of functional mitochondrial DNA during oogenesis restricts the transmission of a deleterious mitochondrial variant. Nat Genet 46, 389–392 (2014).

21. T. Lieber, S. P. Jeedigunta, J. M. Palozzi, R. Lehmann, T. R. Hurd,Mitochondrial fragmentation drives selective removal of deleterious mtDNA in the germline. Nature 570, 380–384 (2019).

22. N. P. Kandul, T. Zhang, B. A. Hay, M. Guo, Selective removal of deletion-bearing mitochondrial DNA in heteroplasmic Drosophila. Nat Commun 7, 13100 (2016).

23. C. L. Samstag et al., Deleterious mitochondrial DNA point mutations are overrepresented in Drosophila expressing a proofreading-defective DNA polymerase gamma. PLoS Genet 14, e1007805 (2018).

24. M. A. Walker et al., Purifying Selection against Pathogenic Mitochondrial DNA in Human T Cells. N Engl J Med 383, 1556–1563 (2020).

25. H. Ma, P.H. O’Farrell, Selfish drive can trump function when animal mitochondrial genomes compete. Nat Genet 48, 798–802 (2016).

26. Z. Chen et al., Genetic mosaic analysis of a deleterious mitochondrial DNA mutation in Drosophila reveals novel aspects of mitochondrial regulation and function. Mol Biol Cell 26, 674–684 (2015).

27. A. C. Chiang, E. McCartney, P. H. O’Farrell, H. Ma, A Genome-wide Screen Reveals that Reducing Mitochondrial DNA Polymerase Can Promote Elimination of Deleterious Mitochondrial Mutations. Curr Biol 29, 4330–4336 e4333 (2019).

28. K. E. White, D. M. Humphrey, F. Hirth, The dopaminergic system in the aging brain of Drosophila. Front Neurosci 4, 205 (2010).

29. F. S. Carlos O. Miller, Malcolm H. Von Saltza, F. M. Strong, Kinetin, a cell division factor from deoxyribonuceic acid. J. Am. Chem. Soc. 77, 1392–1392 (1955).

30. N. T. Hertz et al., A neo-substrate that amplifies catalytic activity of parkinson’s-disease-related kinase PINK1. Cell 154, 737–747 (2013).

31. L. Osgerby et al., Kinetin Riboside and Its ProTides Activate the Parkinson’s Disease Associated PTEN-Induced Putative Kinase 1 (PINK1) Independent of Mitochondrial Depolarization. J Med Chem 60, 3518–3524 (2017).

32. D. C. Samuels, P. Wonnapinij, L. M. Cree, P. F. Chinnery, Reassessing evidence for a postnatal mitochondrial genetic bottleneck. Nat Genet 42, 471-472; author reply 472-473 (2010).

33. Y. S. Ju et al., Origins and functional consequences of somatic mitochondrial DNA mutations in human cancer. Elife 3 (2014).

34. J. N. Jasmin, C. Zeyl, Rapid evolution of cheating mitochondrial genomes in small yeast populations. Evolution 68, 269–275 (2014).

35. E. Bastiaans et al., Regular bottlenecks and restrictions to somatic fusion prevent the accumulation of mitochondrial defects in Neurospora. Philos Trans R Soc Lond B Biol Sci 369, 20130448 (2014).

36. D. Narendra, A. Tanaka, D. F. Suen, R. J. Youle, Parkin is recruited selectively to impaired mitochondria and promotes their autophagy. J Cell Biol 183, 795–803 (2008).

37. A. Bratic et al., Complementation between polymerase-and exonuclease-deficient mitochondrial DNA polymerase mutants in genomically engineered flies. Nat Commun 6, 8808 (2015).

38. N. Rohland, D. Reich, Cost-effective, high-throughput DNA sequencing libraries for multiplexed target capture. Genome Res 22, 939–946 (2012).

39. A. L. Orr et al., Long-term oral kinetin does not protect against alpha-synuclein-induced neurodegeneration in rodent models of Parkinson’s disease. Neurochem Int 109, 106–116 (2017).

